# Large-Scale Network Embedding Predicts Opposing Functions of Adjacent Regions Within Human Prefrontal Cortex

**DOI:** 10.64898/2026.09.02.748980

**Authors:** Anne Billot, Wendy Sun, Kathryn Rodrigues, Xiangyu Wei, Mark C. Eldaief, Randy L. Buckner

## Abstract

Prefrontal regions are hypothesized to be organized hierarchically in support of cognitive control. Using precision functional MRI in three independent cohorts of intensively scanned participants (N=37), we consistently identified a lateral prefrontal cortex (LPFC) region linked to a canonical control network separate from a nearby rostral LPFC region linked to the action-mode network. Despite their spatial juxtaposition, the two regions were differentially coupled to the caudate and ventral putamen, suggesting they are components of segregated networks. Working memory demands activated the canonical LPFC control region but not the adjacent region. Contrasting go and no-go trials during target detection revealed a robust functional double dissociation during goal-directed behavior. The canonical LPFC control region activated when responses were withheld, while the putamen-coupled LPFC region increased activity during executed responses. These findings demonstrate that adjacent LPFC regions participate in opposing functions predicted by their embedding within distinct parallel large-scale networks.

## Introduction

The prefrontal cortex (PFC) participates in executive function and cognitive control (Milner and Petrides 1984; Goldman-Rakic 1987; Shallice and Burgess 1991; Miller 2000; Koechlin et al. 2003; Gilbert and Burgess 2008; Duncan 2010; Fuster 2015; Badre and Nee 2018; Mansouri et al. 2020; Friedman and Robbins 2022). What remains elusive is how the PFC is organized in support of such functions. One prominent class of models proposes a broad progression from lower-level sensory-motor functions in caudal regions to rostral (anterior) PFC regions that support abstracted task goals (Koechlin et al. 2003; Badre and D’Esposito 2009; see also Christoff and Gabrieli 2000; Wagner et al. 2001; Petrides 2005; Badre 2008). The anatomical basis of hierarchical control is hypothesized to be a series of nested corticostriatal loops with the highest-level loop situated on lateral PFC (LPFC), exerting top-down influence on lower-order premotor and primary motor loops that control actions (Badre and Nee 2018).

Recent precision estimates from within-individual human functional neuroimaging reveal that PFC organization is more complex than typically appreciated. Specifically, while detailed analyses of prefrontal organization confirm a broad gradient that progresses from caudal motor regions to rostral association zones, multiple regional specializations deviate from the expectations of a gradient. Neighboring regions can be specialized for distinct processing functions across multiple zones of PFC. For example, within ventrolateral PFC (VLPFC), domain-flexible regions supporting cognitive control lie adjacent to distinct regions specialized for meaning-based linguistic processing (Fedorenko et al. 2013; Blank and Fedorenko 2017; Fedorenko and Blank 2020). Within dorsolateral PFC (DLPFC), regions supporting cognitive control lie adjacent to regions specialized for spatial processing (DiNicola et al. 2023; Du et al. 2024). Additional anatomical distinctions have emerged seemingly idiosyncratically throughout the PFC (e.g., Michalka et al. 2015; Noyce et al. 2017, 2022; Tobyne et al. 2017; Braga and Buckner 2017; Braga et al. 2019; DiNicola et al. 2020; Du et al. 2024; DiNicola and Buckner 2026; Gratton and Braga 2026). The PFC thus possesses both a broad functional organization across its caudal-to-rostral extent, radiating outwards from motor cortex, as well as a patchwork of specialized interdigitated regions. What might be the basis of these local regional specializations?

Goldman-Rakic (1988) proposed that a core determinant of PFC organization is a series of parallel anatomical circuits that support functional specialization of adjacent regions. By this view, the specializations between juxtaposed PFC regions arise because of their embedding within large-scale distributed networks that are distinct but adjacent to one another. This is an appealing framework because it allows anatomical constraints – specifically the anatomical organization of precision identified networks – to be the foundation for functional understanding (e.g., as applied in practice in Du et al. 2024 and as described theoretically as the inside-out approach in Dosenbach et al. 2025). The broad hypothesis is that the functional differences between adjacent regions of the PFC will be understood by considering the anatomy and functional properties of the distinct brain-wide networks in which they are embedded (see also Buckner and DiNicola 2019; DiNicola and Buckner 2021; Buckner 2026; Gordon et al. 2026).

Here we describe a striking functional specialization within LPFC that arises from application of precision neuroimaging within an anatomically focused (inside-out) framework. Across a series of three independent cohorts, we first identified the anatomical positions of distinct adjacent LPFC regions within the idiosyncratic anatomy of each individual participant using network-based functional connectivity. Then, we prospectively tested the functional response properties of the regions across task contrasts. We discovered that adjacent regions of LPFC, each embedded within distinct brain-wide networks, demonstrate opposing functional response properties during a simple target detection task. These results, in combination with growing evidence that the PFC is more specialized than traditionally considered, encourage a revision to how we frame the organization of the PFC. This includes practical implications for clinical translation, especially as precision-targeted approaches to neuromodulation become widespread.

## Results

### Adjacent LPFC regions participate in distinct distributed cortical networks

Our explorations began by identifying two adjacent LPFC regions that are embedded within distinct cortical networks. The first LPFC region is associated with the Frontoparietal Network-A (FPN-A), a canonical cognitive control network also referred to as the multiple-demand system or executive control network (Duncan and Owen 2000; Dosenbach et al. 2006, 2008; Vincent et al. 2008; Fedorenko et al. 2013). FPN-A contains a large LPFC region that extends ventrally (Du et al. 2024; see also Ladwig et al. 2026). The second LPFC region, found dorsal and rostral to the first, is identified via its coupling to cingulo-opercular cortex and is associated with two distinct networks – the Action-Mode Network^1^ (AMN; Dosenbach et al. 2025) and the Salience Network (SN; Seeley 2019). While the AMN and SN can be distinguished by their distributed anatomical patterns (see Dosenbach et al. 2025), they are tightly interwoven in LPFC as shown in the Supplementary Materials (see also Du et al. 2024; Sun et al. 2025). Given this anatomy, we estimated a single, continuous LPFC region associated with both AMN and SN (the *AMN/SN LPFC region*) and a second LPFC region associated with FPN-A (the *FPN-A LPFC region*).

The left panel of **Figure 1** illustrates the adjacent AMN/SN and FPN-A LPFC regions in three representative participants. All participants from the Discovery dataset are shown in the Supplementary Materials. While the two LPFC regions were in the same relative spatial positions across participants, the regions’ exact sizes and border locations varied from one person to the next. In all cases, there was an LPFC region linked to AMN/SN that extended from the middle frontal gyrus into the superior frontal gyrus, and a separate LPFC region linked to FPN-A that extended ventrally and caudally.

**Figure 1:**
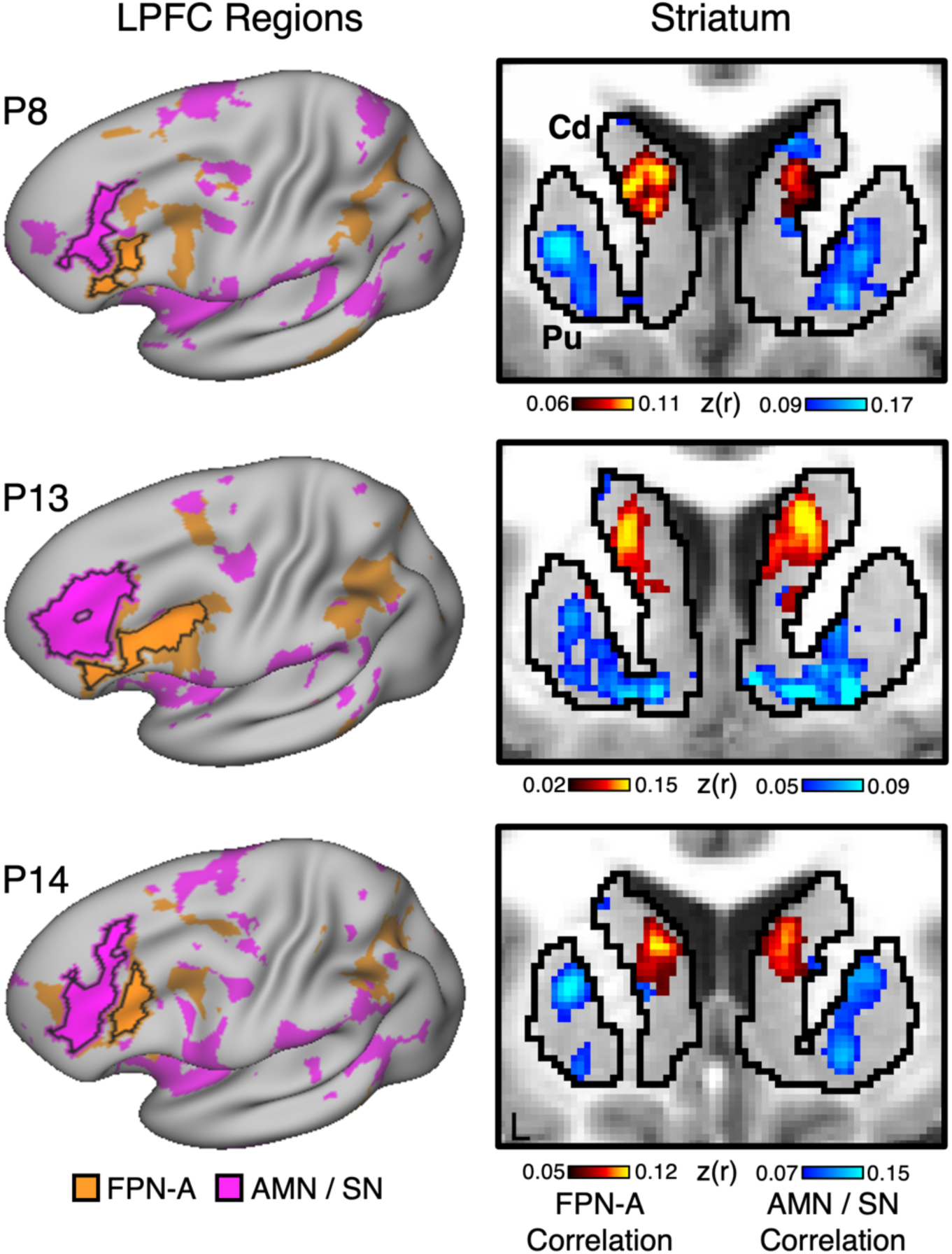
Adjacent lateral prefrontal cortex regions are embedded within parallel distributed cortical networks and coupled to distinct regions of the striatum. The left column displays distinct lateral prefrontal cortex (LPFC) regions associated with the Frontoparietal Network A (FPN-A, orange) and the combined Action-Mode Network/Salience Network (AMN/SN, pink) for three representative individuals from the Discovery dataset (P8, P13, and P14). The regions are visualized on the left cortical surface rotated dorsally. Note that the two LPFC regions are adjacent to one another but coupled to distinct distributed cortical networks. The right column displays the striatal regions correlated with each of the two adjacent LPFC regions. The FPN-A LPFC region is preferentially coupled to the caudate (Cd), whereas the AMN/SN LPFC region is preferentially coupled to the putamen (Pu) extending into ventral striatum. The color scales represent the strength of Fisher *r*-to-*z* transformed Pearson’s correlations with the FPN-A (warm colors) and AMN/SN (cool colors) LPFC regions.

The identification of the adjacent regions was not dependent on the use of the Multi-Session Hierarchical Bayesian Model (MS-HBM). The adjacent LPFC regions could also be revealed by examining seed-region- based correlations that make no assumptions about network organization and do not rely on model priors (see Supplementary Materials). Thus, across individuals and analytic approaches, we consistently identified two separate regions within LPFC that were associated with distinct distributed cortical networks.

### Adjacent LPFC regions link to anatomically distinct zones of the striatum

Further evidence for anatomical separation between the adjacent LPFC regions came from examining correlations with subcortical structures. The cerebral cortex projects to the striatum with partial segregation into cognitive, motor, and affective zones (Heimer 1983; Alexander et al. 1986; Haber 2003). The right panel of **Figure 1** displays the correlation patterns between the adjacent LPFC regions and distinct regions of the striatum. Correlation maps for all participants from the Discovery dataset are shown in the Supplementary Materials.

The adjacent regions of LPFC are associated with anatomically distant zones of the striatum. The LPFC region associated with FPN-A was linked to the caudate, whereas the LPFC region associated with AMN/SN was linked to the ventral putamen. These two zones of the striatum are typically framed as being anatomically and functionally distinct: the FPN-A LPFC region is associated with the prototypical ‘cognitive’ subdivision of the caudate, and the AMN/SN LPFC region is associated with portions of the striatum that fall between the affective (reward/motivation) subdivision (the nucleus accumbens) and the body-mapped motor effector subdivision of the putamen (see also Kosakowski et al. 2024, 2025).

All analyses thus converge to support that the adjacent zones of LPFC are linked to anatomically distinct brain-wide networks, including coupling to different zones of the striatum.

### Adjacent LPFC regions differentially respond to working memory and target detection

We next tested whether the juxtaposed LPFC regions have distinct functional properties by contrasting tasks placing differential demands on working memory and goal-directed action. For each participant, we quantified the functional response level in each of the two LPFC regions during performance of an N-Back working memory task and during target detection in a visual oddball task. Response level estimates were extracted from the individually defined LPFC regions (one value per participant per task) and then pooled at the group level. The results are shown in **Figure 2**.

**Figure 2:**
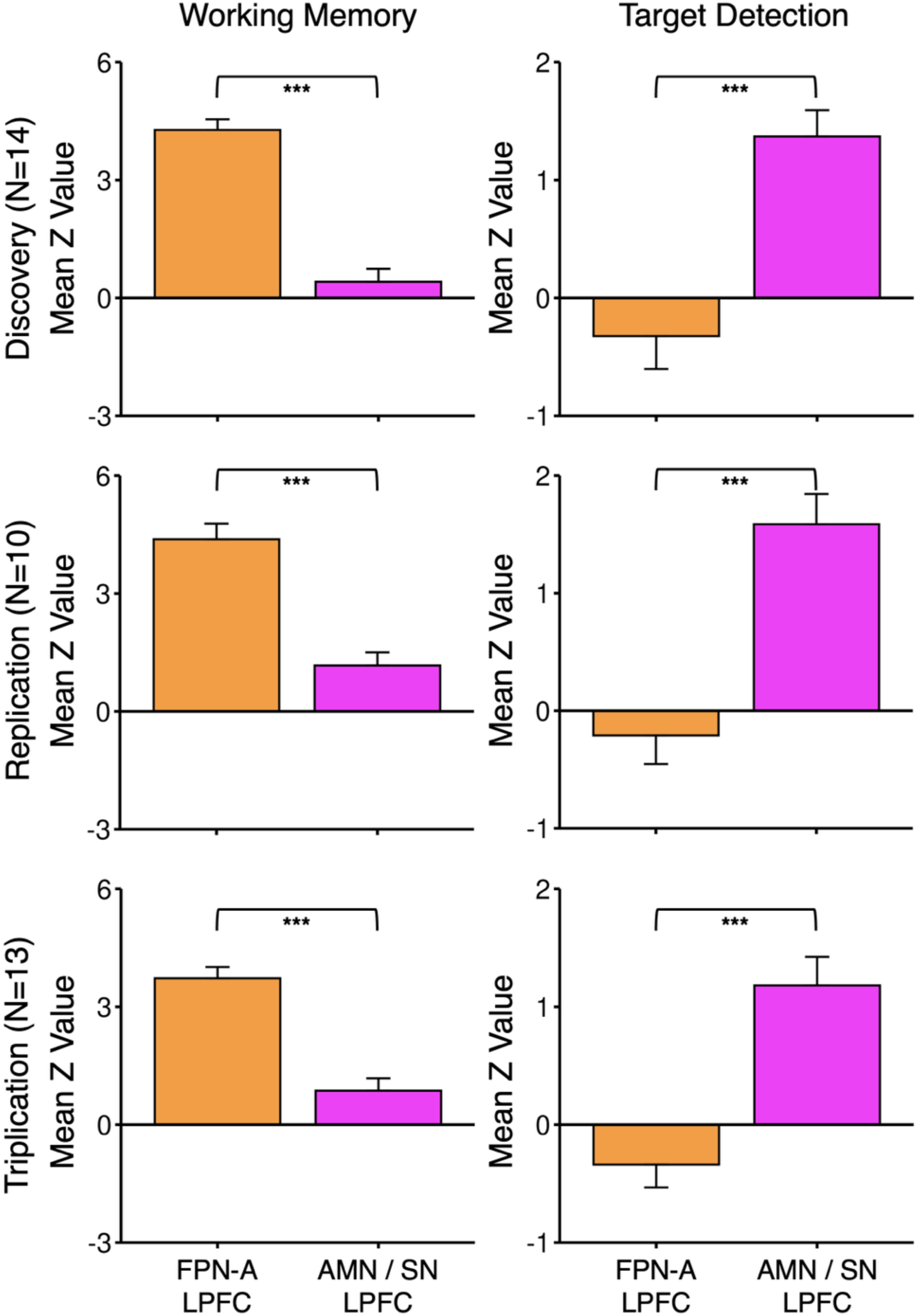
The adjacent lateral prefrontal cortex regions show a robust functional double dissociation. Each plot displays the mean response for the distinct lateral prefrontal cortex (LPFC) regions associated with the Frontoparietal Network-A (FPN-A, orange) and the combined Action-Mode Network/Salience Network (AMN/SN, pink). Error bars show standard error of the mean. The three rows display independent data from the Discovery, Replication and Triplication datasets. The FPN-A LPFC region increases activity during the working memory (N-Back) task. In contrast, the AMN/SN LPFC region increases activity when participants respond during oddball target detection. The interaction between LPFC Region and Task Contrast was significant in all three datasets (*ps* < 0.001) with post-hoc tests establishing the full crossover interaction. *** = *p* < 0.001.

The adjacent LPFC regions displayed a robust functional double dissociation. The FPN-A LPFC region responded during the working memory task but showed minimal response when responding to targets in the oddball task. By contrast, the AMN/SN LPFC region responded when targets were detected but minimally during the working memory task. The interaction between LPFC region (FPN-A versus AMN/SN) and the task (working memory versus target detection) was significant (*F*(1,13) = 121.74, *p* < 0.001, *_η_*^2^*_G_* = 0.66). Post-hoc analyses further showed that the response to the working memory task was significantly higher in the FPN-A LPFC region compared to the AMN/SN LPFC region (*t*(13) = 10.87, *p_adj_* < 0.001, *d* = 2.91, 95% CI [1.67,4.12]); the reverse pattern was found for target detection in the oddball task (*t*(13) = 5.69, *p_adj_* < 0.001, *d* = 1.52, 95% CI [0.73,2.29]).

### Adjacent LPFC regions can be identified and functionally dissociated fully within individuals

We replicated the functional double dissociation in 10 new participants. For these analyses, the LPFC regions were estimated using the same procedures as in the Discovery dataset but, here, task functional response levels were extracted across 10 prospective MRI sessions per participant, allowing the double dissociation to be statistically tested fully within each individual. The Replication dataset also included participants with Major Depressive Disorder (MDD).

Results are illustrated from three healthy participants (**Figure 3**) and three participants with MDD (**Figure 4**). All participants from the Replication dataset are shown in the Supplementary Materials. The interaction between the two LPFC regions and task (working memory versus target detection) was significant in each of the 10 participants (all *ps* < 0.001). Post-hoc analyses further revealed that the response to the working memory task was significantly higher in the FPN-A LPFC region compared to the AMN/SN LPFC region in all 10 participants (all *ps_adj_* < 0.001); the reverse pattern was significant for target detection in the oddball task in all 10 participants as well (all *ps_adj_* < 0.05). See Supplementary Materials for individual-level statistical results.

**Figure 3:**
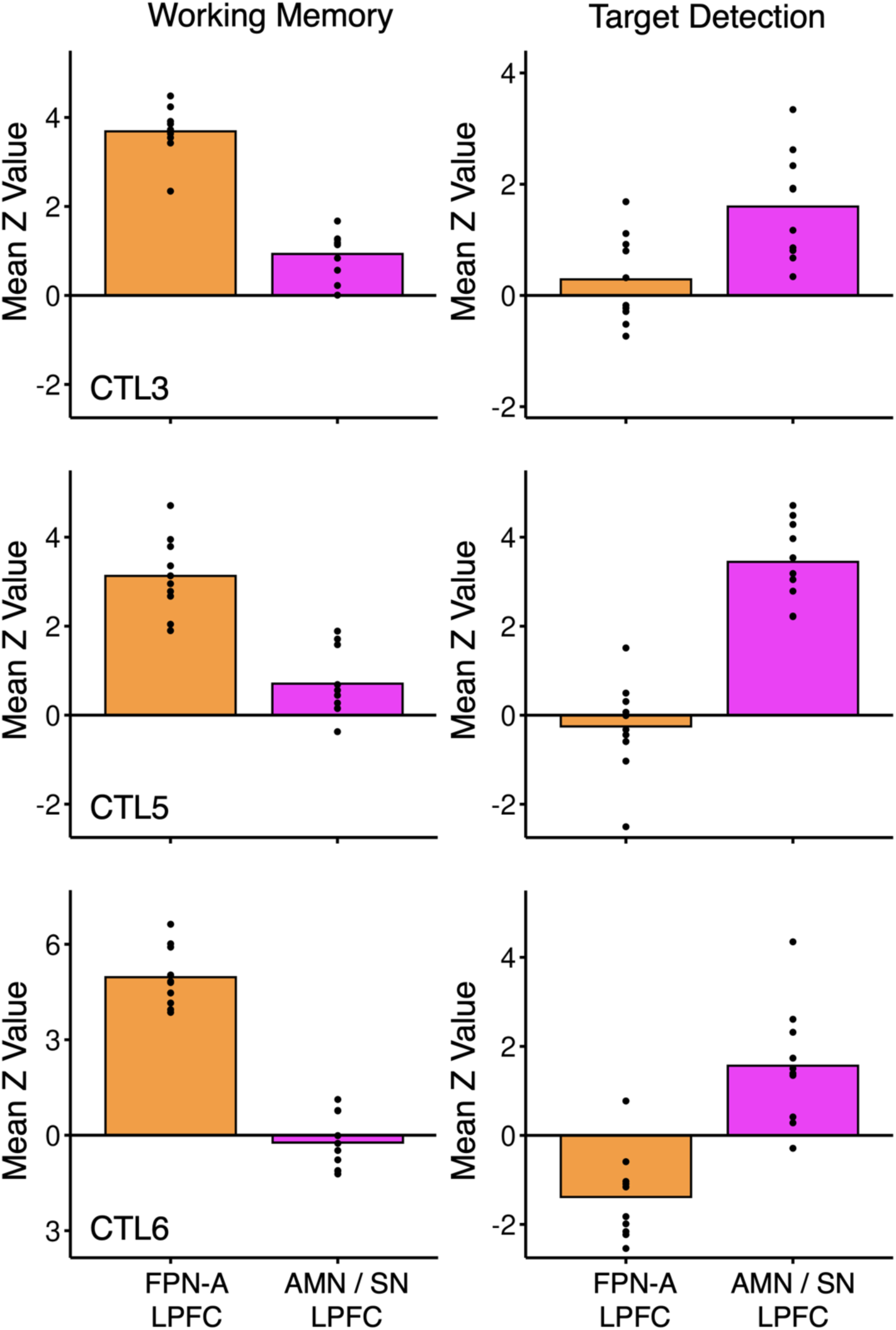
Within-individual functional dissociation. Data from three control participants from the Replication dataset are presented. For each individual, functional networks were first estimated from a single MRI session and then responses were extracted during the working memory (left column) and oddball target detection (right column) tasks from 10 prospective, independent MRI sessions. Bars show the mean response for each participant separately for the adjacent lateral prefrontal cortex (LPFC) regions with data points displaying the response for individual MRI sessions. In each control participant, the FPN-A LPFC region shows a strong, preferential response to the working memory (N-Back) task and a minimal or negative response to the oddball target detection task, while the AMN/SN LPFC region shows the opposite pattern. Data from all control participants are available in the Supplementary Materials.

**Figure 4:**
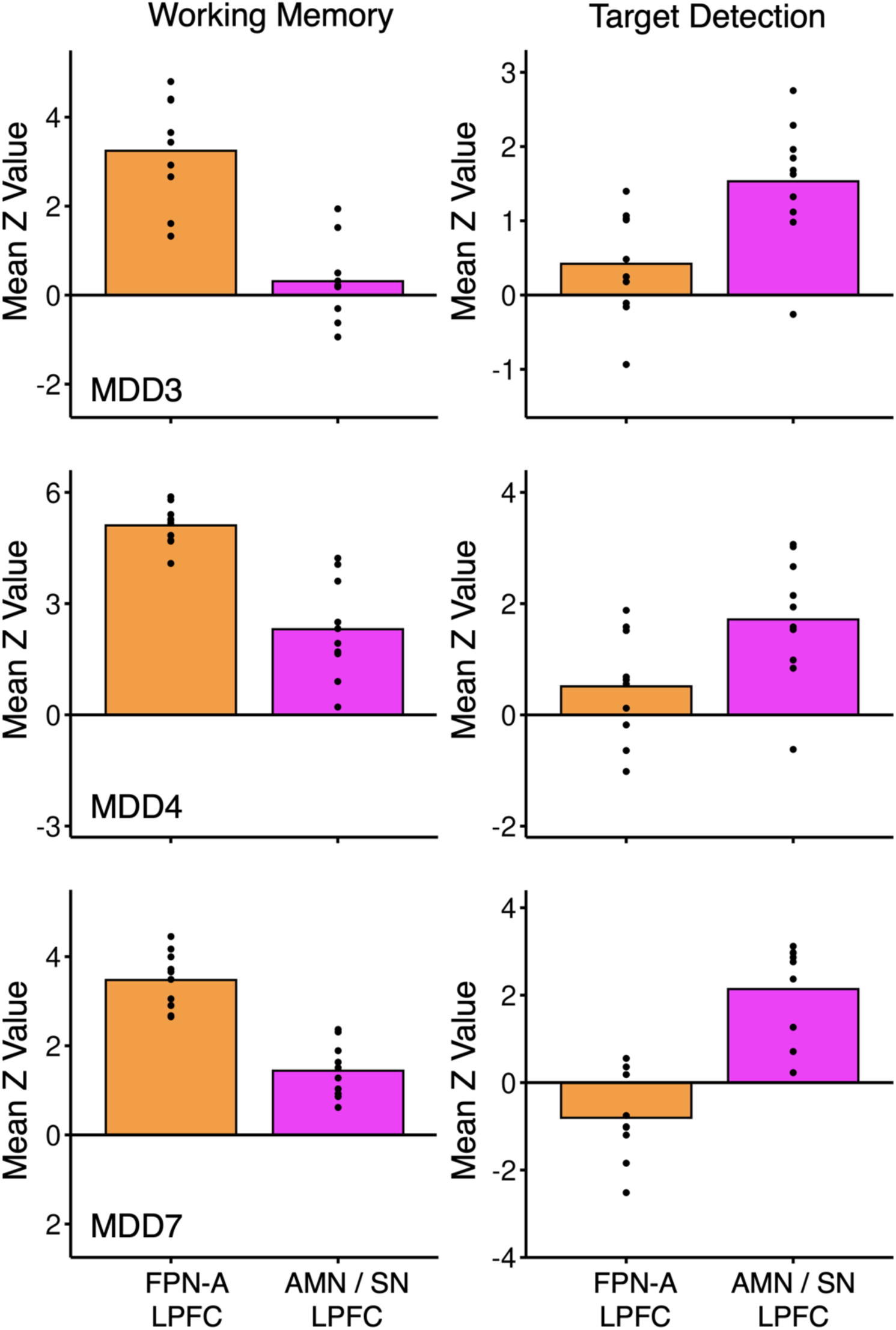
Within-individual functional dissociation generalizes to patients with major depressive disorder (MDD). Data from three participants with MDD from the Replication dataset are presented similar to Figure 3. Bars show the mean response for each participant separately for the adjacent lateral prefrontal cortex (LPFC) regions with individual data points displaying the response for individual MRI sessions. In each participant, the FPN-A LPFC region shows a stronger response to the working memory (N-Back) task and a minimal or negative response to the oddball target detection task, while the AMN/SN LPFC region shows the opposite pattern. Data from all participants with MDD are available in the Supplementary Materials.

When the data from the Replication dataset participants were pooled, paralleling the group-level analysis of the Discovery dataset, the same robust functional double dissociation was again found (see middle row in **Figure 2**). The interaction between the two LPFC regions and the task (working memory versus target detection) was significant (*F*(1,9) = 97.05, *p* < 0.001, *_η_*^2^*_G_* = 0.64). Post-hoc analyses further showed that the response during the working memory task was significantly higher in the FPN-A LPFC region compared to the AMN/SN LPFC region (*t*(9) = 9.12, *p_adj_* < 0.001, *d* = 2.88, 95% CI [1.42,4.32]), and the reverse pattern was observed for target detection in the oddball task (*t*(9) = 5.33, *p_adj_* < 0.001, *d* = 1.69, 95% CI [0.68,2.65]).

### Adjacent LPFC regions can be functionally dissociated by examining no-go trials

To further explore the functional properties of the adjacent LPFC regions, we separately examined the target and salient no-go lure trials within the oddball task. Examining the no-go lure trials is particularly informative because cognitive control demands are high but the motor action must be withheld. Behavioral results indicated that participants made occasional false alarms to the lure trials (1.7% and 1.3% in the Discovery and Replication datasets, respectively) and missed few targets (3.4% and 5.3%).

Examining the functional data revealed another robust double dissociation, in this instance anchored on distinct trial types within the same task. The interaction between the two LPFC regions and trial type (target versus no-go lure) was significant for the Discovery (*F*(1,13) = 37.90, *p* < 0.001, *_η_*^2^*_G_* = 0.32) and Replication (*F*(1,9) = 22.14, *p* = 0.001, *_η_*^2^*_G_* = 0.41) datasets. Post-hoc t-tests indicated the FPN-A LPFC region was more active for the no-go lure trials than the target trials (Discovery dataset *t*(13) = 3.53, *p_adj_* = 0.002, *d* = 0.94, 95% CI [0.30,1.57]; Replication dataset *t*(13) = 1.96, *p_adj_* = 0.041, *d* = 0.62, 95% CI [-0.08,1.29]^2^). The AMN/SN region showed the opposite pattern, being more active for target trials than for no-go lure trials (Discovery dataset *t*(13) = 4.65, *p_adj_* < 0.001, *d* = 1.24, 95% CI [0.52,1.93]; Replication dataset *t*(9) = 4.74, *p_adj_* = 0.001, *d* = 1.50, 95% CI [0.56,2.40]).

To directly visualize the functional response patterns for the two trial types, the time courses were extracted from the individually defined LPFC regions, time-locked to the trial onset for all target trials and for all no-go lure trials in each participant. The time courses were then averaged separately for the Discovery and Replication datasets without any model assumptions. **Figure 5** displays the results. No-go lure trials elicited a robust, transient response in the FPN-A LPFC region, with minimal response in the adjacent AMN/SN region. By contrast, the target trials showed the reverse pattern: the FPN-A LPFC region displayed minimal response, while there was a transient response in the adjacent AMN/SN region.

**Figure 5:**
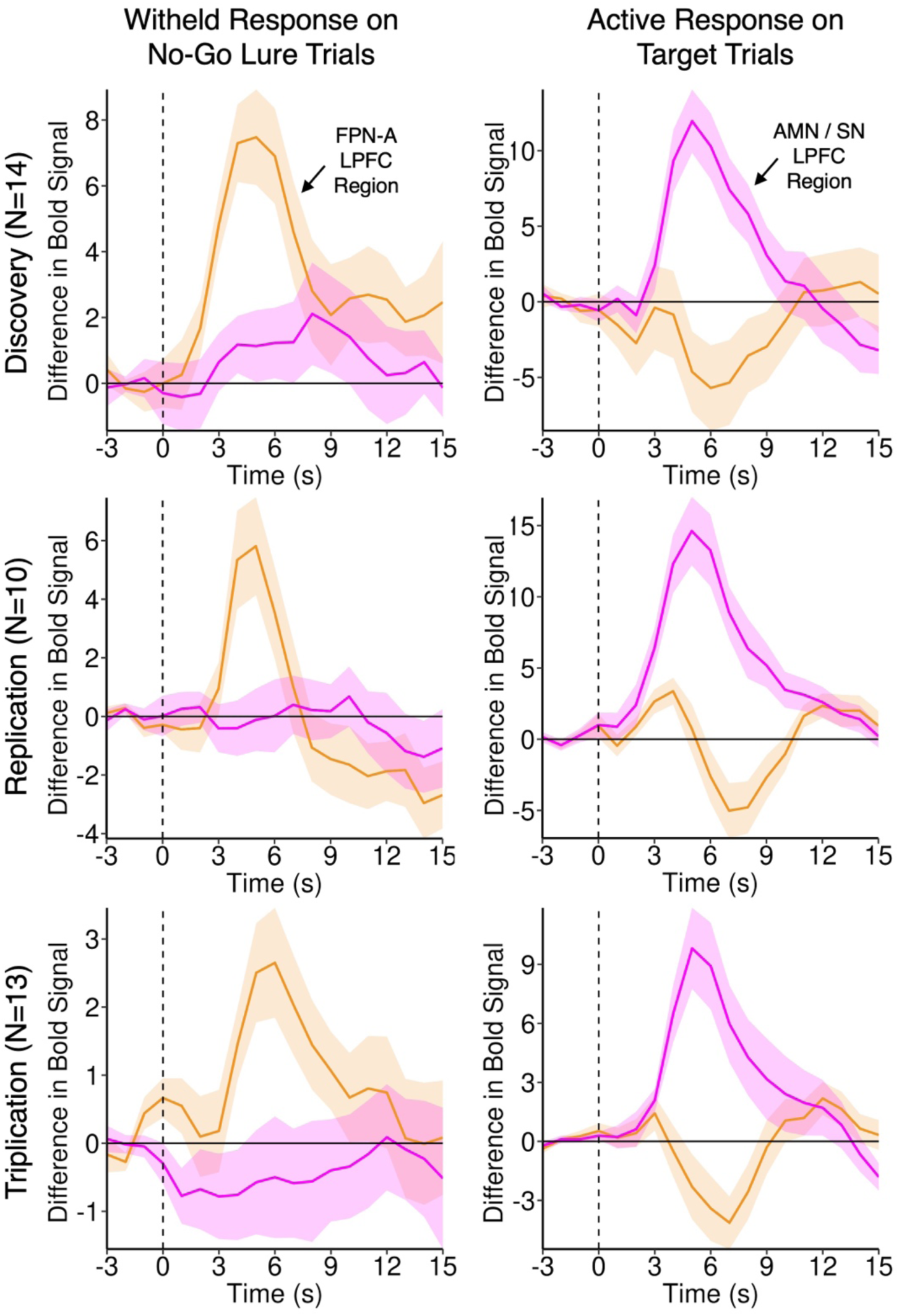
Opposing functional response properties during target detection dissociate the adjacent lateral prefrontal cortex regions. Each time course displays the mean response during the oddball task for the distinct, adjacent lateral prefrontal cortex (LPFC) regions associated with the Frontoparietal Network-A (FPN-A, orange) and the combined Action-Mode Network/Salience Network (AMN/SN, pink). The solid lines show the mean across participants; shading shows the standard error of the mean. The time courses are separated by trial type in the two columns (Left, no-go lure trials; Right, target trials). The three rows display independent data from the Discovery, Replication and Triplication datasets. A robust, transient response in the FPN-A LPFC region is found when responses are withheld (left column), whereas target trials elicit the opposite pattern with a robust, transient response in the adjacent AMN/SN LPFC region (right column).

**Figure 6** displays the mean group activation patterns during working memory, no-go lure trials, and target trials. Across both the Discovery and Replication datasets, working memory and lure trials engage a similar LPFC region situated within the FPN-A network boundaries from the DU15NET-Consensus atlas (Du et al. 2024). By contrast, target trials recruit a distinct, adjacent LPFC region located rostral and dorsal to the FPN-A LPFC region.

**Figure 6:**
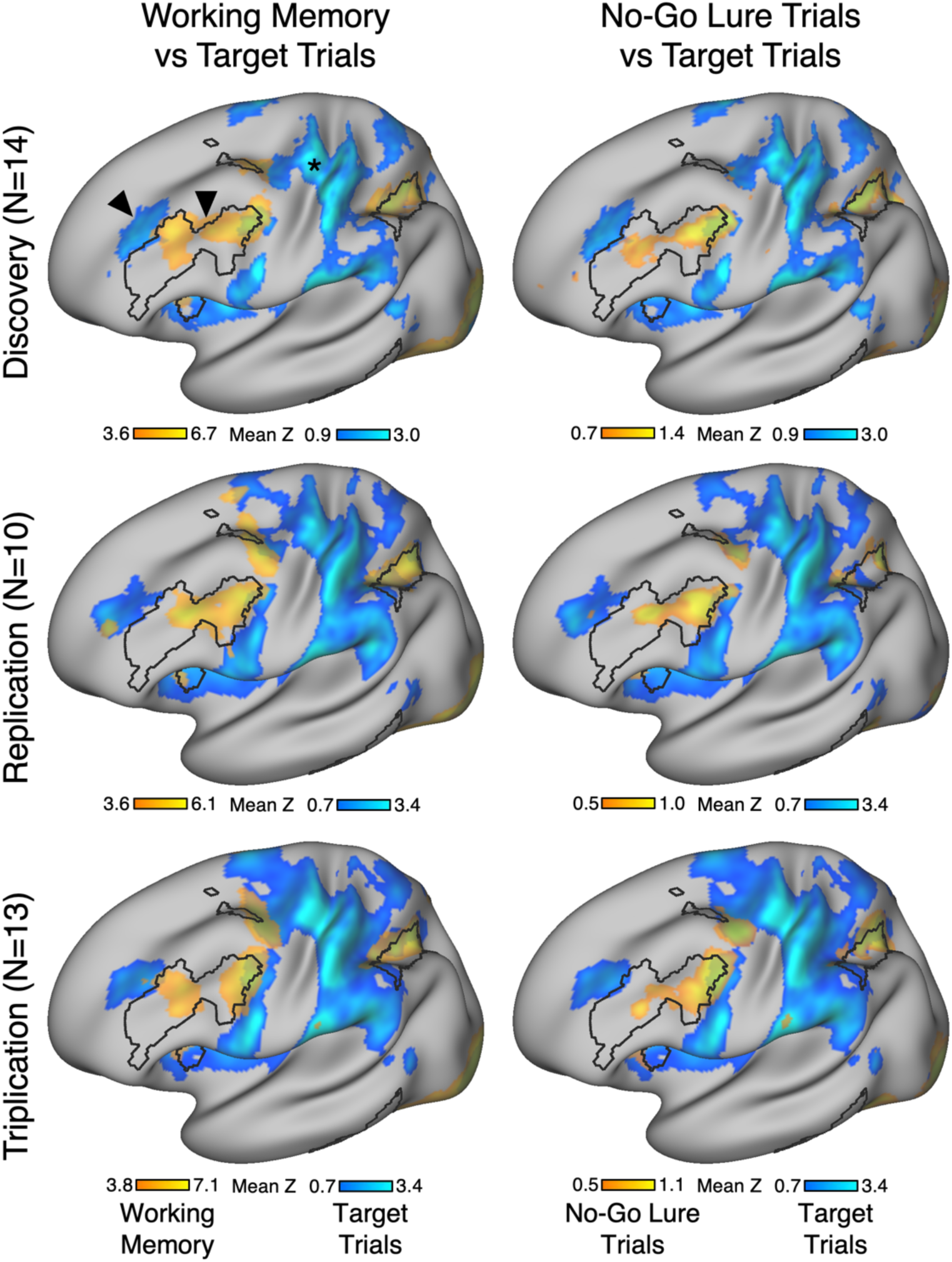
The dissociated lateral prefrontal cortex regions are revealed and replicated across the task maps. Having established a robust functional dissociation between distinct lateral prefrontal cortex (LPFC) regions in targeted analyses, the spatial juxtaposition could be revealed by mapping the critical task contrasts. The left column displays task contrast maps for the working memory (N-Back) task (orange- yellow) simultaneously with the target trial effect (blue) from the oddball task. The right column displays the task contrast maps for the withheld responses during the no-go lure trials (orange-yellow) and the target trial effect (blue) both from the oddball task. The three rows display independent data from the Discovery, Replication and Triplication datasets. The maps show the mean *z* values at the group level. The black outline represents the canonical Frontoparietal Network-A (FPN-A) from DU15NET-Consensus parcellation (Du et al. 2024). Note that the task contrasts consistently reveal distinct LPFC regions in spatially adjacent regions (marked by arrows in the top left panel). Note also the motor cortex response (marked by an asterisk) consistent with the right-handed button press response for target trials.

### Adjacent LPFC regions can be identified and functionally dissociated in a triplication cohort

As a final analysis, we prospectively replicated all key findings in an independent Triplication dataset. Other than the participants being new, analyses were identical to the Replication dataset. Each participant was scanned during a baseline MRI session (from which individually defined LPFC regions were extracted) and then within-individual task responses were measured across 10 prospective MRI sessions.

The interaction between the two LPFC regions and task (working memory versus target detection) was again significant in each of the 13 participants (all *ps* < 0.05, see Supplementary Materials). Post-hoc analyses further revealed that the response to the working memory task was significantly higher in the FPN-A LPFC region compared to the AMN/SN LPFC region in 12 of 13 participants (all *ps_adj_* < 0.001); the reverse pattern was significant for target detection in the oddball task in 10 of 13 participants (all *ps_adj_* < 0.05).

The functional double dissociation was robust when the Triplication dataset participants were analyzed at the group level (see bottom row in **Figure 2**). The interaction between the two LPFC regions and task (working memory versus target detection) was confirmed (*F*(1,12) = 231.40, *p* < 0.001, *_η_*^2^*_G_* = 0.59), with the working memory task response significantly higher in the FPN-A LPFC region compared to the AMN/SN LPFC region (*t*(12) = 7.75, *p_adj_* < 0.001, *d* = 2.15, 95% CI[1.13,3.14]) and the reverse pattern present for target detection in the oddball task (*t*(12) = 5.03, *p_adj_* < 0.001, *d* = 1.40, 95 % CI[0.61,2.16]).

Within the oddball task, the interaction between the two LPFC regions and the trial type (target versus no-go lure) was significant (*F*(1,12) = 32.29, *p* < 0.001, *_η_*^2^*_G_* = 0.35). The FPN-A LPFC region was more active for the no-go lure trials than the target trials (*t*(12) = 3.84, *p_adj_* = 0.001, *d* = 1.06, 95% CI[0.63,1.74]) and the AMN/SN region displayed the opposite pattern (*t*(12) = 4.60, *p_adj_* < 0.001, *d* = 1.28, 95% CI[0.52,2.00]). The bottom row of **Figure 5** shows the time courses for the target and no-go lure trials.

Working memory and no-go lure trials activated the same LPFC region at the group level and this region aligns with the FPN-A network boundaries in the DU15NET-Consensus atlas (Du et al. 2024). In contrast, target trials activated a separate LPFC cluster located rostral and dorsal to this FPN-A LPFC region (see bottom row of **Figure 6**).

## Discussion

Human prefrontal cortex exhibits marked specialization. Using precision neuroimaging, we discovered and replicated a robust functional dissociation between nearby LPFC regions that, despite their close proximity, participate in distinct distributed cortical networks, display differential coupling to the caudate and putamen, and exhibit opposing response properties. One LPFC region, falling within the canonical cognitive control network, was active during working memory and transiently engaged when motor actions had to be withheld during target detection – a pattern consistent with a role in cognitive control that extends to response suppression. A juxtaposed LPFC region showed the opposite pattern: it was minimally engaged during working memory but responded robustly when targets elicited an action, consistent with its role in the execution of goal-directed behavior as a component region within the AMN. These collective findings suggest LPFC comprises multiple differentially specialized regions whose properties arise from the large-scale networks in which they are embedded.

### Prefrontal specialization arises from regional embedding in segregated brain-wide networks

Architectonic features, differences in extrinsic connectivity patterns, and the differential behavioral consequences of brain lesions have established that primate prefrontal cortex possesses specialized subregions (Sanides 1969; Goldman-Rakic 1987; Barbas and Pandya 1989; Petrides and Pandya 1994; Öngür and Price 2000; Stuss and Alexander 2007; Passingham and Wise 2012; Fuster 2015). Human within-individual precision neuroimaging studies have found multiple functional dissociations between adjacent regions in prefrontal cortex (e.g., Fedorenko et al. 2012; Michalka et al. 2015; Noyce et al. 2017; DiNicola et al. 2023; Du et al. 2024; Deen and Freiwald 2025; DiNicola and Buckner 2026; Ladwig et al. 2026). A major open question is how these subregions are organized, and what underlying principles explain why each is specialized in the way that it is.

Drawing on primate anatomical connectivity, Goldman-Rakic (1988) proposed that prefrontal specialization, including functional distinctions between adjacent areas, can be understood by examining the large-scale networks in which the adjacent regions are embedded (see Buckner 2026 for discussion). The present results in the human can be conceptualized within such a framework. The two LPFC regions dissociated here were identified via their participation in distinct large-scale networks. The first region was embedded within FPN-A. FPN-A is a distributed frontoparietal network that participates in working memory and other forms of task that place demands on cognitive control (variably known as the “multiple-demand”, “frontoparietal control”, or “executive control” network; Duncan and Owen 2000; Dosenbach et al. 2008; Vincent et al. 2008; Fedorenko et al. 2013). The second region was embedded within a composite of the AMN and SN. Of most direct interest is the AMN^3^, a distributed network that includes regions within the dorsal anterior cingulate cortex and frontal operculum, that is active during the execution of goal-directed behavior (Dosenbach et al. 2006, 2025).

When the two LPFC regions were isolated within individuals, a robust functional double dissociation emerged that was reliable across independent datasets and task contrasts. Specifically, the LPFC region associated with FPN-A strongly activated during the N-Back working memory task (**Figure 2**). This result is predicted by group-level investigations that have consistently found increases along the inferior frontal gyrus as working memory load increases (e.g., Braver et al. 1997; Jonides et al. 1997; Wager and Smith 2003; Owen et al. 2005). Using intensive repeated sampling, we further demonstrated that the effect is detectable within individuals (**Figures 3-4**). Examining the same region during target detection revealed that it also responded robustly when participants had to withhold their response – a role consistent with response inhibition or a related aspect of cognitive control (**Figure 5**). Critically, the adjacent LPFC region, linked to the AMN, responded preferentially when salient targets were acted upon, but not during working memory or when responses were withheld (**Figure 5**), echoing distinctions between LPFC regions implicated in response execution as contrast to response inhibition (Cieslik et al. 2013).

Taken together, these results demonstrate that nearby regions of LPFC participate in distinct brain-wide networks with markedly different functional properties. A parsimonious possibility is that, as the cortex develops from primary sensorimotor through intermediate to tertiary association cortex, these territories fractionate and specialize, with adjacent subregions inheriting the processing specializations of the networks in which they are embedded. This developmental cascade may give rise to a complex mosaic of specialized regions throughout the cerebral cortex, prominently including PFC (Du et al. 2024; Saadon-Grosman et al. 2024; Buckner 2026; Gordon et al. 2026).

### Limitations and future directions

The present analyses reveal novel spatial details about PFC organization. The methods are nonetheless limited by resolution. For example, a combined AMN/SN region was used in the present analyses (similar to Sun et al. 2025). AMN and SN are spatially adjacent, sometimes even interdigitated, and thus are difficult to separate especially within LPFC. Results for each network considered separately are presented in the Supplementary Materials revealing convergent response patterns. This result might be a property of spatial limitations or alternatively a point of functional convergence between the two networks under these specific task conditions. Future work at higher spatial resolution using elaborated task designs will be required to provide further insight.

Second, the present evidence is limited in its temporal resolution. The hemodynamic response sampled by fMRI cannot resolve the millisecond-scale dynamics by which the two regions are recruited within a single trial. Temporal dynamics may constrain mechanistic hypotheses. Intracranial recordings in neurosurgical patients alongside precision fMRI offer a particularly promising avenue for this question, as they combine the spatial precision needed to dissociate adjacent LPFC regions with the temporal resolution needed to characterize their relative dynamics during target detection and response inhibition.

The present findings may be informative for therapeutic neuromodulation, particularly for repetitive transcranial magnetic stimulation (rTMS) in the treatment of depression. In prior work, we demonstrated that standard-of-care approaches, including scalp-based anatomical targeting and functional-connectivity-based targeting, target different functional networks across individuals. Through modeling, we found that the recently FDA-approved rTMS protocol targeting the LPFC engages an AMN/SN region in most individuals but an FPN-A region in others (Sun et al. 2025). The opposing functional response patterns of these two regions suggest that they may have distinct therapeutic effects, consistent with evidence that distinct LPFC networks map onto dissociable symptoms (Siddiqi et al. 2020, 2022; Tozzi et al. 2024). Specifically, the FPN-A region may contribute to cognitive symptoms such as difficulty concentrating and emotion regulation deficits (Kaiser et al. 2015; Hack et al. 2023), and the AMN/SN region may contribute to anhedonia and motivational symptoms (Borsini et al. 2020; Pizzagalli and Roberts 2022). While the present study does not address therapeutic mechanisms, the discovery that side-by-side LPFC regions can be embedded within distinct distributed networks and show opposing functional properties motivates further study of whether targeting of one region versus the other yields differential therapeutic benefit, and for whom.

## Conclusions

The functional organization of human PFC is characterized by a patchwork of specialized regions whose functions are predicted by the distributed networks in which they are embedded. Two such regions sit side by side in LPFC and, despite their anatomical proximity, are coupled to distinct striatal regions and support opposing functions: one tracks cognitive control demands during working memory and when action must be withheld, while the other responds when goal-directed action is initiated. This double dissociation was replicated in two independent cohorts and could be observed within individual participants. These findings underscore the value of within-individual precision neuroimaging for resolving PFC organization.

## Methods

### Discovery dataset

We first identified and tested functional response properties in an initial dataset – the Discovery dataset, and then subsequently replicated the findings in two additional datasets as described below. The Discovery dataset was previously reported in Du et al. (2024) and includes participants who completed intensive repeated MRI sampling; methodological details are briefly repeated here. For the present paper, individualized functional networks were estimated from resting-state fMRI data and used to define adjacent LPFC regions. Correlations were examined between the LPFC regions and the striatum within individuals to establish that the regions are components of separate brain-wide networks. Task fMRI data were then used to evaluate response properties of these PFC regions.

#### Participants

Fourteen healthy right-handed English-speaking adults contributed data to the Discovery dataset (age range: 18-34 years old, mean = 22.0, SD = 4.1, 9 female). One original participant from Du et al. (2024) was excluded due to an insufficient number of usable oddball task runs. History of a neurological or psychiatric illness was an exclusion criterion. Participants provided informed consent using procedures approved by the Institutional Review Board of Harvard University.

#### MRI Data Acquisition

Data were acquired at the Harvard Center for Brain Science with a 3-T Prisma^fit^ MRI scanner using a 32-channel phased-array head coil (Siemens Healthineers, Erlangen, Germany). Participants were instructed to remain still and alert and to look at a rear-projected display through a mirror attached to the head coil. Foam and inflated padding mitigated head motion. During scans, eyes were monitored for alertness using an EyeLink 1000 Plus with Long-Range Mount (SR Research, Ottawa, ON, Canada), and motion was monitored using Framewise Integrated Real-time MRI Monitoring (FIRMM; Touring Medical, St. Louis, MO, USA).

High-resolution T1-weighted (T1w) and T2-weighted (T2w) structural images were acquired using Human Connectome Project (HCP) sequences (Harms et al. 2018). T1w MPRAGE parameters: voxel size = 0.8 mm, TR = 2,500 ms, TE = 1.81, 3.60, 5.39, and 7.18 ms, TI = 1,000 ms, flip angle = 8°, matrix 300 x 320 x 208, 208 slices, in-plane GRAPPA acceleration = 2. T2w SPACE parameters: voxel size = 0.8 mm, TR = 3,200 ms, TE = 564 ms, matrix = 300 x 320 x 208, 208 slices, in-plane GRAPPA acceleration = 2. As backup, T1w structural scans were also collected using a multi-echo MPRAGE sequence (van der Kouwe et al. 2008): voxel size = 1.2 mm, TR = 2,200 ms, TE = 1.57, 3.39, 5.21, and 7.03 ms, TI = 1,100 ms, flip angle = 7°, matrix 192 x 192 x 144, 144 slices, in-plane GRAPPA acceleration = 4.

fMRI data were acquired using a multiband gradient-echo echo-planar pulse sequence provided by the University of Minnesota sensitive to blood oxygenation level-dependent (BOLD) contrast (e.g., Feinberg et al. 2010; Xu et al. 2013): voxel size = 2.4 mm, TR = 1,000 ms, TE = 33.0 ms, flip-angle = 64°, matrix 92 x 92 x 65 (FOV = 221 x 221), 65 slices, anterior-to-posterior (AP) phase encoding, multislice 5x acceleration.

Resting-state fixation runs were collected during which participants fixated on a central black crosshair. Each run was 7 min 2 s, with 422 frames; the first 12 frames were removed for T1 equilibration. Participants completed 17-24 resting-state fixation runs (∼2 per session) and 48 to 70 task fMRI runs across 8-11 sessions. To mitigate spatial distortions, dual-gradient-echo B0 field maps were acquired at each session (TE = 4.45, 6.91 ms with slices matched to the BOLD sequence).

Each BOLD fMRI run was examined for quality. Exclusion criteria used parameters reported in Xue et al. (2021), including 1) maximum absolute motion >1.8 mm and 2) slice-based SNR <130. Runs with SNR >100 but also SNR <130 were retained if motion and visual inspection indicated adequate quality. All fMRI data exclusions were finalized before network estimation.

#### MRI Data Preprocessing and Network Identification

Data were preprocessed with an openly available pipeline (“iProc”), which minimizes spatial blur and preserves individual idiosyncratic anatomy with a single interpolation step for BOLD data (see detailed description in Braga et al. 2019), using tools from FreeSurfer (Fischl 2012), FSL (Jenkinson et al. 2012), and AFNI (Cox 2012). Preprocessed data were taken directly from Du et al. (2024). The MS-HBM was implemented to estimate cortical networks within individuals (Kong et al. 2019; Du et al. 2024). First, the connectivity profile of each vertex on the fsaverage6 cortical surface was estimated as its functional connectivity to 1,175 regions of interest (ROIs) uniformly distributed across the fsaverage surface (Yeo et al. 2011). For each run of data, the Pearson’s correlation coefficients between the fMRI time series at each vertex (40,962 vertices/hemisphere) and the 1,175 ROIs were computed. The resulting 40,962 × 1,175 correlation matrix was then binarized by keeping the top 10% of the correlations to obtain the functional connectivity profiles (Yeo et al. 2011). Next, the expectation-maximization (EM) algorithm for estimating parameters in the MS-HBM was initialized with a group-level parcellation from a subset of the HCP S900 data release (that itself used the clustering algorithm from our previous study; Yeo et al. 2011). The 15 networks are labeled Somatomotor-A (SMOT-A), Somatomotor-B (SMOT-B), Premotor-Posterior Parietal Rostral (PM-PPr), Action-Mode (AMN), Salience (SN), Dorsal Attention-A (dATN-A), Dorsal Attention-B (dATN-B), Frontoparietal Network-A (FPN-A), Frontoparietal Network-B (FPN-B), Default Network-A (DN-A), Default Network-B (DN-B), Language (LANG), Visual Central (VIS-C), Visual Peripheral (VIS-P), and Auditory (AUD). To confirm that the individual network estimates were not obligated by the assumptions and ensure that network estimates properly captured individual correlation patterns, a seed region-based analysis was conducted as described in Du et al. (2024).

#### Construction of Prefrontal Regions

After networks were estimated within individuals, two adjacent LPFC regions within FPN-A and AMN/SN were identified within each hemisphere. These LPFC regions consisted of the largest contiguous cluster of the FPN-A and AMN/SN within a prefrontal search space in fsaverage6 space at the surface level (Lynch et al. 2022). If the two largest clusters from the respective networks were not adjacent, the second largest cluster of one of the networks identified as the closest to the largest cluster of the other network was selected through visual inspection. The resulting clusters were then eroded by 1 mm on the inflated pial surface with the Connectome workbench command -metric-erode version 1.3.2 (Marcus et al. 2013) to account for some level of unavoidable spatial blurring. If the erosion divided the region into multiple clusters, only the largest ones were kept after visual inspection. The selection process of example LPFC regions in three participants in the left hemisphere are displayed in Supplementary Materials and all other LPFC regions are also shown in Supplementary Materials. All LPFC regions were defined before any further analyses were performed to ensure an unbiased estimate of their properties.

#### Within-Individual Striatum-to-Cortex Correlation Matrices

The methods to identify the striatum and compute striatum-to-cortex correlation matrices within individual participants were adapted from Kosakowski et al. (2024, 2025). To isolate striatal voxels, we identified the caudate, putamen, and nucleus accumbens in each participant using the FreeSurfer automated parcellation (Fischl et al. 2002). The FreeSurfer-based masks were combined, binarized, and warped using the same individual-specific analysis pipeline used in “iProc” preprocessing (Braga et al. 2019).

Both resting-state fixation and task fMRI runs were used to estimate the striatum-to-cortex correlation matrix within individual participants. Task fMRI runs used the same preprocessing pipeline as the resting-state fMRI runs, focusing on the residuals after removing contributions of the trial and block task effects via regression (Du et al. 2025; see also Fair et al. 2007). We included residualized data from task runs because it substantially increases the available data to improve signal-to-noise properties, which benefits estimation of subcortical correlation patterns (yielding 47-94 total runs of data for each participant in the Discovery dataset). Specifically, we ran a task fMRI general linear model (GLM) analysis (details described for each task in Du et al. 2024) and included the resulting task-regressed data (residuals) in constructing the correlation matrix in the same manner as for resting-state data (see Du et al. 2025 for validation of the approach).

For each participant, the pairwise Pearson’s correlation coefficients between the fMRI time courses at each surface cortical vertex were calculated for each run, yielding an 81,924 x 81,924 matrix (40,962 vertices/hemisphere). For the subcortex, we used a volume mask including the striatum. For each run, we computed pairwise Pearson’s correlation coefficients for the fMRI time courses between each volume voxel within the mask and each cortical vertex. The subcortical matrix was 134,797 x 134,797 and the subcortical to cortical matrix was 134,797 x 81,924. The matrices were then Fisher *r*-to-*z* transformed and averaged across all runs to yield a single best estimate of the within-individual correlation matrices. These *z*-scored matrices were combined and assigned to a cortical and subcortical template, combining the left and right hemispheres of the fsaverage6 surface and subcortex of the MNI152 volume into the CIFTI format to interactively explore correlation maps using the Connectome Workbench’s wb_view software (Marcus et al. 2013). For each LPFC region, we averaged the vertex-wise correlations between all vertices in the region and every striatum voxel, yielding a single vector describing the region’s average correlation with the entire striatum.

#### Oddball Task Paradigm

The oddball task explored detection of transient responses to salient, visual oddball targets that were uncommon relative to irrelevant and distracting nontargets (similar to Wynn et al. 2015). The goal of the task was to activate the AMN/SN. Both the AMN and SN have regions at or near the anterior insula and have been variably associated with response to task-relevant transients (see Seeley et al. 2007; Seeley 2019 and Dosenbach et al. 2006, 2025 for discussion).

Participants viewed a sequence of uppercase letters O and K in either black or red. Participants pressed a button with their right index finger when a red K appeared and withheld their responses to all the other letter-color combinations. The random trial ordering was set with OptSeq (Dale 1999). In each run, 10% of the trials were target red Ks, 10% were no-go lure red Os, 40% were distractor black Ks, and 40% were distractor black Os. The two contrasts of interest were either the target red Ks or the no-go lure red Os versus all other non-target trials coded as the implicit baseline.

Each run lasted 5 min 50 s (350 frames, with the first 6 frames removed for T1 equilibration). After 6 s of fixation overlapping the initial stabilization frames, an initial 20-s block of fixation was followed by a continuous extended block of 300 1-s trials (0.15 s presentation of the letter followed by 0.85 s of fixation) and then a final 20-s block of extended fixation. Before the first trial a 2-s start cue (1-s “Begin,” 1-s fixation) was presented, as well as a similar “End” cue after the final trial. Thus, the design was a rapid, event-related paradigm sandwiched between blocks of extended fixation. Five runs were collected for each participant. Runs were excluded from the analysis if participants missed more than six targets within a task run, which accounted for 20% of the total targets. Usable oddball task runs included in the analysis ranged from 3 to 5. Our primary contrasts examined responses to target and no-go lure trials against the implicit baseline corresponding to non-target trials.

#### Working Memory (N-Back) Task Paradigm

The working memory (N-Back) task was extended from Braver et al. (1997) to explore demands on cognitive control under varied levels of memory load. Specifically, the N-Back task included a 2-Back versus 0-Back comparison to target FPN-A. However, in this study, we only used the 2-Back versus Fixation contrast to allow for qualitative comparisons of the results between the three datasets. Stimuli were presented sequentially in the center of the computer screen. Participants maintained fixation on a central crosshair throughout the run.

The stimuli varied across four conditions (Face, Word, Scene, and Letter) that were each presented in separate blocks. Faces and scenes were color images, with scenes showing both indoor and outdoor spaces and chosen not to feature people (faces from HCP, Barch et al. 2013; scenes from Konkle and Oliva 2012, Josephs and Konkle 2020). Letters included subsets of consonants, and words featured one-syllable words from 10-word sets matched for length and frequency with the Corpus of Contemporary English (Davies 2010, December 2015 version). In all but the Word condition, participants matched the stimuli to an exact stimulus referent (0-Back), or the exact stimulus presented two trials before (2-Back). For the Word condition, the participants decided whether the current word rhymed with the target (e.g., “dream” would be a positive match with “steam”).

Each N-Back run featured eight blocks (a 0-Back and a 2-Back for each of the 4 stimulus categories). Each block included a cue and nine trials. During the first cue stimulus, participants also saw the block type, either 2-Back or 0-Back. The background was black (matching the HCP format). All blocks included two target and two lure (repeated nontarget) trials. Targets and lures were equally likely to appear in each viable trial position within and across runs. Participants pressed a button for every trial, indicating match (right index finger) or no match (left index finger).

Each run lasted 4 min 44 s (284 frames, with the first 12 frames removed for T1 equilibration). After 12 s of fixation overlapping the initial stabilization frames, an additional block of 12 s of fixation was followed by two 25-s blocks of the N-Back task interspersed with 15-s fixation blocks. Across runs, 0-Back and 2-Back blocks, categories, and their interactions were counterbalanced. Each trial was 2.5 s in duration (2 s of stimulus presentation followed by 0.5 s of fixation). The fixation crosshair was white for the extended fixation blocks and green during the N-Back task blocks. Within a run, all categories were seen before a category repeated. Eight runs were collected for each participant. Runs where participants missed responses in more than two trials were excluded from analysis. Usable N-Back task runs included in the analysis ranged from 6 to 8. All task fMRI exclusions were performed before completing any task analysis. Our primary contrast examined the working memory load effect (2-Back vs. fixation) across all stimulus categories.

#### Task Analysis

Functional task data were analyzed with the general linear model (GLM) as implemented by FSL’s first-level FEAT (FSL version 5.0.4; Woolrich et al. 2001). The data were only whole-brain-signal regressed and high-pass filtered with a cutoff of 100 s (0.01 Hz) to remove low-frequency noise within each run. GLM outputs included, for each contrast, β values for each vertex that were converted, within FEAT, to *z* values. Within each participant, *z*-value maps from all runs were averaged together by using fslmaths (Smith et al. 2004) to create a single cross-session map for each contrast of interest on the surface. For the N-Back task, we ran block-level GLMs. The GLM outputs included z-value maps for each block, which were averaged by condition across runs.

To further investigate the BOLD response independent of the GLM design at the onset of oddball task trials, mean BOLD signal from high-pass filtered data was averaged across vertices within each PFC region at each TR. Event-related segments (3 TRs pre-, 15 TRs post-trial onset) were extracted for target and no-go lure trials and averaged within each run, then across runs within individuals, and finally across individuals for group-level plots. These time courses were then plotted as mean signal intensity (± standard error of the mean) across TRs, baseline-corrected to the mean of the 3 TRs pre-trial onset.

#### Statistical Analysis

For each participant and each task contrast, the mean *z* value was calculated across vertices of the individualized bilateral FPN-A and AMN/SN LPFC regions. We hypothesized two functional double dissociations. First, we predicted a dissociation between the N-Back working memory and the oddball target detection tasks, with the FPN-A LPFC region responding preferentially to the working memory task and the AMN/SN LPFC region responding preferentially to the oddball target detection task. Second, we predicted a dissociation within the oddball task itself: the FPN-A LPFC region would respond more strongly to no-go lure trials than to target trials, reflecting cognitive control demands associated with response inhibition, while the AMN/SN LPFC region would show the opposite pattern given the role of the AMN in goal-directed actions. These two hypotheses were tested separately each using a two-way repeated-measures ANOVA with Task Contrast (two levels) and LPFC region (FPN-A versus AMN/SN) as within-subject factors and mean *z* value as the dependent variable using the *rstatix* package in R v4.4.2. Next, we performed post-hoc pairwise comparisons to evaluate differences in mean *z* value between adjacent LPFC regions in response to each task contrast using one-tailed paired t-tests. P-values were Holm-corrected for multiple comparisons and effect sizes are reported as Cohen’s d_z_ (Lakens 2013). 95% confidence intervals for d_z_ were derived from the non-central t distribution (Cumming 2012), implemented in the *effectsize* R package.

### Replication and Triplication datasets

Findings from the Discovery dataset motivated the analyses performed in the Replication and Triplication datasets, which focused on replicating the functional double dissociations at the individual-subject level as well as the group level. In these datasets, we estimated networks from a single MRI session and tested whether these individually defined networks could predict distinct task-evoked responses across an independent set of 10 task fMRI sessions collected with each participant. Thus, this was a stringent test of the double dissociation. MRI acquisition and processing pipelines, task design and statistical analyses followed similar procedures to those described for the Discovery dataset, with differences outlined below.

#### Participants

Right-handed English-speaking adults with major depressive disorder (MDD) and controls were recruited to participate for payment in a study investigating the effects of individualized TMS targeting (Sun et al. 2025). The data were analyzed here combining across MRI sessions independent of the TMS manipulation to replicate the dissociations observed in the Discovery dataset. After exclusion based on behavioral performance (see criteria in the *Task Paradigms* section), the Replication dataset included 7 participants with MDD (age range: 24-39, mean = 32.6, SD = 5.9; 4 female) and 3 controls (CTL, age range: 24-34, mean = 28.7, SD = 5.0; 1 female) and the Triplication dataset included 6 participants with MDD (age range: 20-45, mean = 35.3, SD = 10.1, 3 female) and 7 controls (age range: 22-47, mean = 29.3, SD = 8.5, 3 female). Participants were from diverse racial and ethnic backgrounds (8 of the 23 individuals self-reported as non-white or Hispanic). Exclusion criteria included history of bipolar disorder, schizophrenia, or neurological disorder for all participants, and any psychiatric disorder for controls. Participants provided informed consent using procedures approved by the Institutional Review Board of Massachusetts General Hospital.

#### MRI Data Acquisition

Data were acquired similarly to the Discovery dataset, except for the differences described below. Each participant participated in an initial baseline MRI session and then 10 additional MRI sessions spread over 5 days, distributed over up to 3 months except for one control participant (CTL3) who completed the study within 7 months. High-resolution T1-weighted (T1w) and T2-weighted (T2w) structural images were acquired. T1w MPRAGE parameters: voxel size = 1.0 mm, TR = 2,530 ms, TE = 1.69, 3.55, 5.41, and 7.27 ms, TI = 1,100 ms, flip angle = 7°, matrix 256 x 256 x 192, 192 slices, in-plane GRAPPA acceleration factor = 2. T2w SPACE parameters: voxel size = 1.0 mm, TR = 3,200 ms, TE = 564 ms, matrix = 256 x 256 x 192, 192 slices, in-plane GRAPPA acceleration = 2. Backup T1w structural scans were again collected. Due to artifacts in acquisition for one participant, the backup 1.2mm T1w image was used for preprocessing.

fMRI data were acquired using the same sequence as the Discovery dataset. Each participant completed 8 resting-state fixation runs during the baseline session, with a 10-15 min break in the middle of the session. During each of the subsequent 10 fMRI sessions, they completed one resting-state fixation run, two N-Back working memory task runs, and two oddball task runs. The resting-state run always occurred second, the order of the other two tasks was counterbalanced across sessions within a day, and no task was repeated across two successive runs. One of the MDD participants (MDD2) only completed 8 fMRI sessions after the baseline. To mitigate spatial distortions, three dual-gradient-echo B0 field maps were acquired during the baseline session and two during each of the 10 subsequent fMRI sessions (TE = 4.45, 6.91 ms with slices matched to the BOLD sequence). MRI data quality was monitored during each MRI session using ScanBuddy (Asay et al. 2025). Each BOLD fMRI run was examined for quality using exclusion criteria as in the Discovery dataset. Eight N-Back task runs were excluded due to software inconsistencies in the frame rate of the MRI screen display. Usable resting-state runs for the baseline session ranged from 6 to 8 runs. Usable task runs across the 10 subsequent fMRI sessions ranged from 13 to 20 for the N-Back task and from 8 to 20 for the oddball task after exclusion for poor MRI data quality or task performance (see criteria in *Task Paradigm*s section). All fMRI data exclusions were again finalized before network estimation.

#### MRI Data Preprocessing, Network Identification, Region Construction, and Task Analysis

MRI data from the baseline session and from the 10 subsequent MRI sessions were preprocessed separately using the same “iProc” pipeline as in the Discovery dataset, with the exception that a more expansive motion scrubbing procedure was adopted given the more variable data quality (the present data were from community samples, in contrast to the Discovery dataset which had extremely low motion given recruitment from a college sample). Specifically, after the single interpolation step, confounding variables including 6 head motion parameters, whole-brain signal, ventricular signal, deep cerebral white matter signal, and temporal derivatives as well as the quadratic term were calculated from the data (36 parameters; Ciric et al. 2017). In addition, volumes with high framewise displacement (defined as >0.4mm or >3 standard deviations above the mean of the runs) were flagged in each run and added to the 36-parameter nuisance regression matrix. These signals were regressed out from all BOLD runs (AFNI 3dTproject). Then, only resting-state data were bandpass filtered at 0.01–0.1 Hz (AFNI 3dBandpass). Finally, all BOLD runs were projected to the fsaverage6 cortical surface mesh using trilinear interpolation and smoothed using a 4-mm full-width at half-maximum Gaussian kernel along the cortical surface (FreeSurfer mri_vol2surf and mri_surf2surf).

Cortical networks were estimated within individuals following the exact same approach as in the Discovery dataset but using resting-state fixation data from the single baseline MRI session (i.e., up to 8 runs). The same procedure as in the Discovery dataset was followed for the selection of prefrontal FPN-A and AMN/SN LPFC regions. LPFC regions in all participants of the Replication and Triplication datasets are displayed in Supplementary Materials. The reason the networks and derived LPFC regions were estimated from the baseline session was that the subsequent (independent) sessions were set aside. In this manner, for every individual, the LPFC regions could be defined and then tested for task effects in entirely independent within-subject data.

#### Oddball and Working Memory (N-Back) Task Paradigms

The task designs were similar to the Discovery dataset with additional practice and some modification to simplify the working memory (N-Back) task. Each participant practiced online at home through the Pavlovia platform (https://pavlovia.org/) and task performance was reviewed before the first MRI session. If scores were low, the participant completed additional practice. In addition, at the beginning of each set of two MRI sessions, participants practiced the N-Back task again on a computer outside of the scanner. Practice runs followed the same design as the fMRI runs, using different stimulus sets. Up to 20 runs were collected per task for each participant across 10 sessions (two per session). The oddball task design was identical to the one used for the Discovery dataset, except that a new set of randomized stimuli was created for each run with OptSeq following the same rules as in the Discovery dataset (Dale 1999) and the background was off-white. Runs were excluded from the analysis if participants missed more than 35% of the total targets and participants were excluded from the analyses if the average number of missed targets across all runs was greater than 25%. No participant was excluded in this study based on this criterion.

The working memory (N-Back) task design was simplified to include only the 2-Back and fixation blocks. The stimuli were also simplified to use two conditions (Face, Word). The background was off-white. Faces were color images generated with StyleGAN (Karras et al. 2019), with an equal representation of male and female faces and diversity across racial phenotypes. Words featured one-syllable words from 10-word sets matched for length and frequency with the SUBTLEXUS corpus (Brysbaert and New 2009).

Each N-Back run featured four blocks (two blocks for each of the two stimulus categories). Each block included a cue and nine trials. During the first cue stimulus, participants saw the type of task (2-Back) and the first stimuli (Face or Word). All blocks included two target and two lure (repeated nontarget) trials. Targets and lures were equally likely to appear in each viable trial position within and across runs. Participants pressed a button for every trial, indicating match (right index finger) or no match (left index finger). Each run lasted 3 min 27 s (207 frames, with the first 12 frames removed for T1 equilibration). After 12 s of fixation overlapping the initial stabilization frames, an additional block of 15 s of fixation was followed by 30-s blocks of the N-Back task interspersed with 15-s fixation blocks. Target and lure trials order was counterbalanced across runs and experimental days. Category orders were set for each day, with equal numbers of Face and Word blocks in each block position, and the same category order maintained between experimental days. Each trial was 3 s in duration (2.5 s of stimulus presentation followed by 0.5 s of fixation). The fixation crosshair was black for the extended fixation blocks and green during the N-Back task blocks. Runs where participants missed responses in more than 35% of match trials were excluded from the analysis and participants were excluded if the average number of missed match trials across all runs was greater than 25%. Three control participants were excluded based on this criterion. All task fMRI exclusions were performed before completing any task analysis.

Tasks analyses replicated those used for the Discovery dataset, including the same statistical procedures at both the individual-level response as well as group-level analyses. The regional measures, time courses, and group-level statistical results thus represent direct prospective replications.

## Software and Code Availability

Functional connectivity was computed as Pearson product-moment correlations in MATLAB (v2019a for the Discovery dataset and v2019b for the Replication and Triplication datasets; MathWorks, Natick, MA). Image preprocessing was carried out using a combination of FreeSurfer v6.0.0, FSL, and AFNI. Cortical surface maps were generated in Connectome Workbench v1.3.2, which was also used for model-free seed region verification. Statistical analyses were conducted in R v4.4.2.

## Data Availability

Individual participant data are available through the NIH repository (https://nda.nih.gov).

## Acknowledgments

We thank the Harvard Center for Brain Science neuroimaging core and FAS Division of Research Computing for their support. We thank Tim O’Keefe for assistance in optimization of data processing, Elham El Hallak for assistance with data analysis, and Ross Mair for MRI physics support. The multi-band EPI sequence was generously provided by the Center for Magnetic Resonance Research (CMRR) at the University of Minnesota.

## Footnotes

1 The Action-Mode Network (AMN) has historically been referred to as the Cingulo-Opercular Network (as used in Du et al. 2024). We update the name to reflect Dosenbach et al. (2025).

2 P-values reflect one-tailed paired t-tests (pre-specified directional hypotheses); 95% CIs are reported two-sided throughout for interpretability. In this replication, the two-sided interval narrowly includes zero, consistent with the marginal one-tailed p-value.

3 AMN and SN were combined in the present study because of their tight juxtaposition in LPFC. Given the response pattern, we discuss the relations to AMN, and discuss the limitation that future work, possibly at higher resolution using direct neurophysiological recordings, will likely be needed to disambiguate the functional roles of the two regions.

